# Integrated luminescence and phenotypic profiling for drug discovery in a zebrafish model of Marfan syndrome

**DOI:** 10.64898/2026.05.12.722859

**Authors:** Marina Horvat, Lisa Caboor, Karo De Rycke, Lisa Mennens, Elise Daniels, Janne Wyseur, Esther Verhelst, Ilian Roos, Isaac Rodríguez-Rovira, Gustavo Egea, Julie De Backer, Patrick Sips

## Abstract

**Background:** Marfan syndrome (MFS) is a life-threatening heritable connective tissue disorder caused by pathogenic variants in fibrillin-1, characterized by progressive cardiovascular disease. Current medical therapies slow disease progression but do not prevent major complications, underscoring the need for new treatment strategies and unbiased discovery approaches.

**Methods:** We used a zebrafish model of MFS lacking *fibrillin-3* (*fbn3^-/-^*), which recapitulates key cardiovascular phenotypes including cardiac stress, valvular defects, arrhythmia, and aortic dilation. To enable sensitive, quantitative assessment of cardiac stress, we generated a novel transgenic zebrafish reporter expressing secreted nanoluciferase under control of the stress-responsive *nppb* promoter. This reporter was combined with morphological phenotyping and bulbus arteriosus (BA) imaging. We evaluated standard MFS therapies, targeted modulators of TGF-β signaling, and performed an unbiased high-throughput drug screen of over 1 500 clinically approved compounds across multiple developmental treatment windows.

**Results:** *fbn3^-/-^* larvae exhibited markedly elevated *nppb* activity that correlated with phenotypic severity and peaked during stages of highest mortality. The nanoluciferase reporter provided a ∼1 000-fold dynamic range, substantially outperforming Firefly luciferase-based assays. Pharmacological inhibition of TGF-β signaling produced transient or deleterious effects, while β-blockers, losartan, and allopurinol failed to consistently improve cardiac stress, pericardial edema, or BA dilation. The unbiased high-throughput drug screen identified a small number of primary and secondary hits; however, none demonstrated reproducible phenotypic rescue upon rigorous multi-dose, multi-time window validation.

**Conclusions:** This study establishes a sensitive zebrafish-based platform for early, quantitative assessment of cardiovascular stress in MFS. Our findings highlight the limited efficacy of current therapies, the context-dependent nature of TGF-β modulation, and the biological complexity underlying MFS pathogenesis. Although no definitive therapeutic candidates were identified, this work lays a robust foundation for expanded unbiased discovery efforts aimed at identifying disease-modifying interventions for MFS.

## BACKGROUND

Marfan syndrome (MFS) is a major cause of heritable thoracic aortic disease, caused by pathogenic variants in fibrillin-1 (*FBN1*), a large extracellular matrix (ECM) glycoprotein widely distributed throughout the body^1,2^. Structurally, FBN1 acts as a central architectural organizer of the ECM, forming elastic and non-elastic microfibrils. Through their tissue-specific assembly, these microfibrils endow organs with distinct mechanical properties, ranging from skeletal constraint and vascular recoil to lens zonule rigidity and skin pliability^3^. Functionally, FBN1 serves as a critical regulator of extracellular signaling by controlling the bioavailability of transforming growth factor-β (TGF-β) and bone morphogenetic proteins (BMPs), which are essential for tissue development, homeostasis, and repair^4,5^. Consequently, abnormalities in FBN1 give rise to a broad spectrum of clinical manifestations^6^.

The most life-threatening manifestation of MFS is aortic disease. Disruption of aortic ECM homeostasis leads to changes in the aortic wall, including vascular smooth muscle cell (VSMC) phenotypic switching and apoptosis, protease dysregulation, ECM degradation, and intimal injury, which promote thrombosis, chronic inflammation, and impaired wall perfusion. Together, these processes weaken the aortic wall, leading to progressive dilation (aneurysm) and/or predisposing it to dissection and rupture - life-threatening complications that represent the most severe manifestations of aortopathies^7^. Beyond aortic disease, FBN1 deficiency also affects cardiac structure and function. Mitral valve prolapse with regurgitation as well as abnormalities of the aortic valve are common in patients with MFS^8^, while in a subset of patients, biventricular dilated systolic and diastolic dysfunction can be observed^8,9^. Patients are also prone to arrhythmias, including ventricular ectopy and non-sustained ventricular tachycardia, which can lead to sudden cardiac death^10^. In its most severe neonatal form, MFS can cause heart failure and death within months after birth^11^.

Over the years, many different therapeutic avenues have been explored in preclinical studies using MFS mouse models. Ranging from TGF-β-neutralizing antibodies and matrix metalloproteinase inhibitors to modulators of nitric oxide signaling, GABA pathway inhibitors, antioxidants, and even female hormones and chromatin remodeling agents, mouse drug screens have identified a broad spectrum of promising therapeutic targets, as reviewed in Deleeuw *et al^12^*. However, their relative efficacy varies across developmental and disease stages, and translation to clinical trials has not been met with great success.

Current medical therapies are not curative. The standard of care remains centered on antihypertensive agents, such as β-adrenergic receptor blockers (β-blockers) and angiotensin II receptor blockers (ARBs), which reduce the hemodynamic stress exerted on the weakened aortic wall, which can slow, but cannot prevent potentially fatal aortic complications^13^.

This highlights the need for more effective therapies, particularly those that move beyond the limitations of current target-driven approaches. Addressing this gap will require exploration of previously uncharted therapeutic strategies, moving beyond predefined targets toward unbiased discovery approaches. With its small, optically accessible heart and aorta, the zebrafish provides a powerful platform for modeling cardiovascular disease and enabling unbiased high-throughput drug discovery^14^.

Recently, we generated a zebrafish model of MFS, lacking the zebrafish fibrillin-3 gene (*fbn3^-/-^*), which exhibits a consistent, early-onset MFS-like phenotype, including aortic dilation, valvular defects, and arrhythmia^15^. *fbn3^-/-^* larvae demonstrate pericardial edema, which varies in severity, associated with progressive endocardial detachment in the atrium. While all mutants exhibit compromised endocardial integrity, only severely affected embryos develop complete endocardial detachment and die between 6-9 days post-fertilization (dpf) due to obstruction of blood flow, whereas mildly affected mutants can survive to adulthood. *fbn3* mutants also have a dilated bulbus arteriosus (BA), the zebrafish anatomical equivalent of the human aortic root^16,17^ - the primary site of aortic pathology in MFS^18^. These morphological phenotypes of the *fbn3*^-/-^model can be quantified during early developmental stages, making it well-suited for drug screening.

In previous high-throughput screening efforts for cardiovascular disease, cardiac stress in zebrafish embryos was used as a readout. A transgenic reporter line expressing firefly luciferase (Fluc) under control of the *nppb* promoter was developed previously to evaluate hypertrophic cardiomyopathy^19^ and was used successfully in a high-throughput drug screen (HTDS) for arrhythmogenic cardiomyopathy (Naxos disease)^20^. *nppb* encodes a ventricular stress-responsive natriuretic peptide^21^ whose prohormone NT-proBNP is a well-established clinical biomarker of heart failure^22^. Interestingly, circulating levels of NT-proBNP are elevated in MFS patients^23^. We therefore generated an enhanced *nppb*-based reporter using secreted nanoluciferase to enable bioluminescent detection of cardiac stress with a high dynamic range as an additional quantifiable readout in our *fbn3*^-/-^ model.

In this study, we use this novel luciferase transgenic line of an established zebrafish MFS model to assess existing therapies and perform an unbiased HTDS to uncover novel treatment strategies.

## METHODS

### Zebrafish husbandry

Zebrafish were maintained in a semi-closed recirculating housing system (ZebTec, Tecniplast) at a constant temperature (27-28 °C), pH (∼7.5), conductivity (∼550 mS), and a light/dark cycle (14 h/10 h). Fish were fed twice daily with dry food (Gemma Micro, Skretting) and once with Micro Artemia (Ocean Nutrition). Breeding and the collection of the embryos were performed according to previously described protocols^24^. All experiments were performed on offspring from stable *fbn3* mutant lines derived from heterozygotes outcrossed for at least four generations to wild-type (WT) zebrafish to avoid confounding off-target effects.

### nppb:secNluc reporter construction

A three-fragment transgenic construct was generated using NEBuilder® HiFi DNA Assembly (NEB). The vector backbone contained *α-crystalline:yellow fluorescent protein (YFP)*, origin of replication (ORI) and ampicillin resistance (pDest399 pCM326 pdestTol2CY; a kind gift from Prof. MacRae, Brigham and Women’s Hospital, Boston, USA). The *nppb* promoter was amplified from the in-house *Tg(nppb:Fluc)* zebrafish line^19^ (a kind gift from Prof. MacRae), and the coding sequence for secreted nanoluciferase (secNluc) was amplified from pNL1.3-CMV (Promega). Primer pairs are listed in Supplementary Table 1.

DNA fragments were purified using either the QIAquick PCR Purification Kit (Qiagen) or gel extraction kit (QIAquick® Gel Extraction Kit, Qiagen; Zymoclean™ Large Fragment DNA Recovery Kit, Zymo Research). Construct assembly was performed according to the manufacturer’s instructions and transformed into 5-alpha competent *E. coli* (NEB). Positive colonies were screened by restriction digestion with ClaI (NEB) following plasmid isolation using the QIAprep Spin Miniprep Kit (Qiagen). The correct clone was expanded, and plasmid DNA was purified using a PureLink™ HiPure Plasmid Midiprep Kit (Invitrogen, Thermo Fisher Scientific).

### Generation of nppb:secNluc transgenic zebrafish line

The transgenic construct was integrated into zebrafish embryos using the CRISPR/Cas9 system, directed to the safe harbor site *uobl6^25^*. crRNA and tracrRNA (IDT) were annealed at a 1:1 molar ratio to generate gRNA duplexes, which were complexed with Sniper-Cas9 protein (ToolGen). Approximately 200 pg of plasmid construct and Cas9-gRNA ribonucleoprotein complexes were co-injected into one-cell stage WT and *fbn3^-/-^* embryos (FemtoJet, Eppendorf). Transgenic founders were identified by ocular YFP expression.

### Pharmacological treatments of zebrafish embryos

All compounds were dissolved in dimethyl sulfoxide (DMSO) and subsequently diluted in E3 medium to the desired final concentration, with the final DMSO concentration not exceeding 1% (v/v). Solutions were refreshed daily. WT embryos were treated in parallel to evaluate genotype-specific differences in compound toxicity. Highest concentrations used corresponded to the maximal non-toxic doses identified in preliminary screenings.

Transgenic line *Tg(kdrl:GFP)* was used for *in vivo* imaging experiments, whereas *Tg(nppb:Fluc)^19^* and *Tg(nppb:secNluc)* were used for luciferase-based assays. Dose-response experiments were performed for hit validation following the high-throughput drug screen, using a concentration range of 1 nM - 100 µM and n ≤ 18 embryos per condition. For individual dose validation experiments, n = 50-70 embryos per condition were used. In the HTDS, six embryos were allocated per drug (well).

Compounds selected for testing were: refametinib, RepSox, metoprolol succinate, atenolol, nebivolol, losartan, and allopurinol. A diverse chemical library of over 1500 compounds, previously approved or developed for other applications, was used for the HTDS.

### Phenotypic evaluation of drug-treated embryos

Compound-treated *fbn3^-/-^* zebrafish were morphologically evaluated at 3, 4, and 5 dpf, unless stated otherwise. Mortality was noted and surviving larvae were categorized as exhibiting either a severe (S) or mild (M) phenotype based on the size of pericardial edema, with severity defined as pericardial edema extending beyond the ventral edge of the yolk sac (Figure 1D). For each treatment, we quantified the proportion of embryos exhibiting the severe *fbn3*^−/−^ phenotype and compared it with a solvent-treated group.

**Figure 1.**
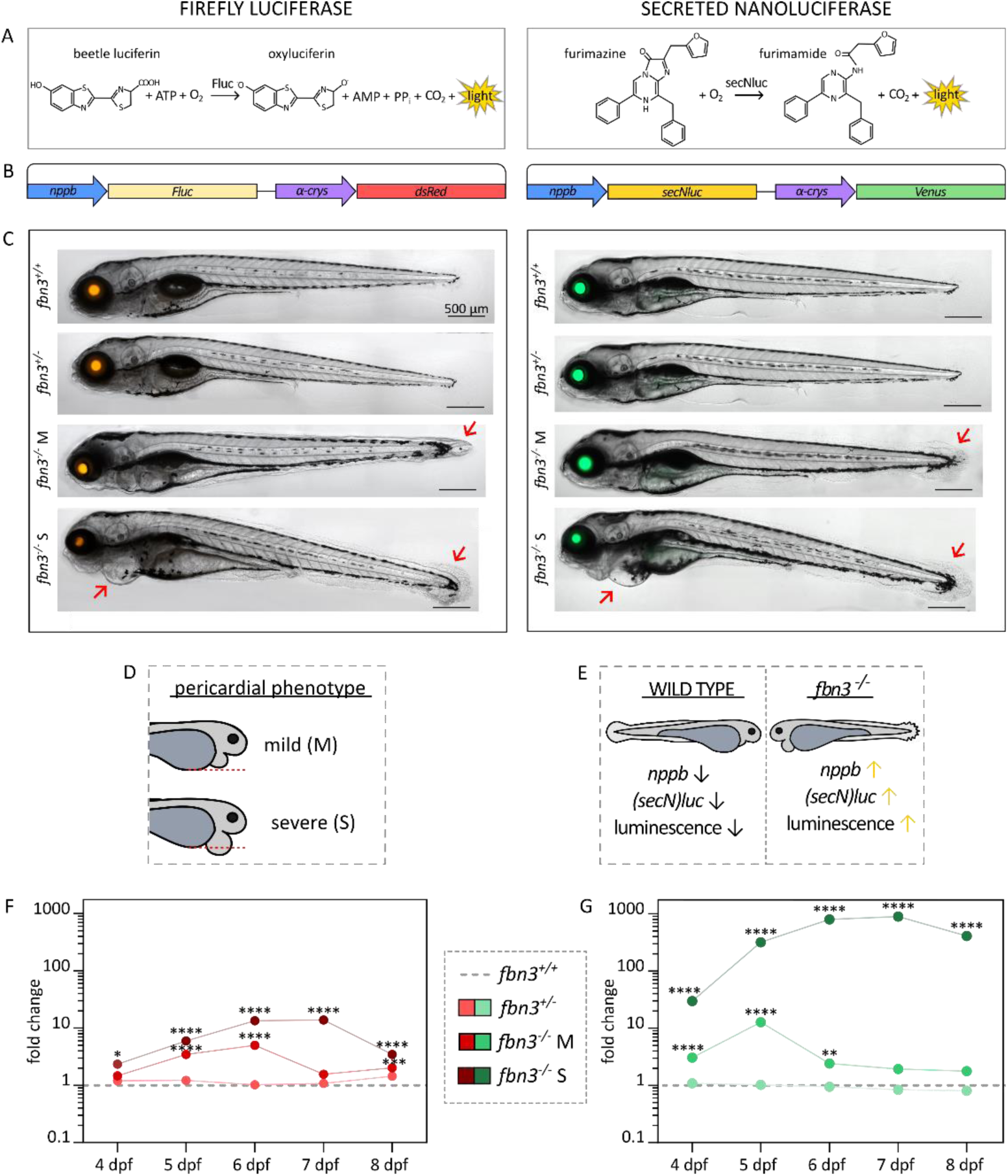
Transgenic bioluminescent reporters for cardiac stress reveal elevated nppb-driven luminescence in fbn3 mutant zebrafish. **(A)** Chemical reactions catalyzed by Firefly luciferase (Fluc) and secreted nanoluciferase (secNluc) reactions. **(B)** Transgene architecture of the luciferase reporter lines. Both luciferases are under the control of the cardiac stress-responsive *nppb* promoter and include an *α-crystallin* (*α-crys*)-driven fluorescent marker (*DsRed* or *Venus*), indicating successful transgene integration. **(C)** Transgenic *nppb*:luciferase zebrafish WT and *fbn3^-/-^* embryos, with fluorescent lens signals (red/orange (*DsRed*) – Fluc, green (*Venus*) – secNluc) indicating transgene integration. Red arrows highlight characteristic *fbn3* mutant phenotypes, including pericardial edema and finfold atrophy. **(D)** *fbn3^-/-^* larvae categorization as severe (S) or mild (M) based on the size of pericardial edema, with severity defined as pericardial edema (red dotted line) extending beyond the ventral edge of the yolk sac. **(E)** Schematic summary of reporter output in WT vs *fbn3*^-/-^ mutants. Loss of *fbn3* function leads to increased *nppb* activity and elevated bioluminescence. **(F, G)** Fold-change in luminescence measured across developmental stages (4-8 dpf) in *fbn3*^+/-^, *fbn3*^-/-^ (mild, M), and *fbn3*^-/-^ (severe, S) larvae, plotted relative to WT controls (grey dotted line). secNluc (G) generates up to 100-fold greater luminescence than Fluc (F), allowing highly sensitive monitoring of *nppb*-dependent stress responses. Data are expressed as mean ± SEM. Statistical test analysis: unpaired t-test on log-transformed data (F, G). ****p<0.0001, ***p<0.001, **p<0.01, *p<0.05. M = mild *fbn3^-/-^*pericardial phenotype, S = severe *fbn3^-/-^* pericardial phenotype.

### In vivo imaging of zebrafish bulbus arteriosus

At 6 or 8 dpf, zebrafish larvae were positioned on their ventral side and embedded in 1% low-melting temperature agarose (SeaPlaque, Lonza). 200-frame-long videos (at 25 frames per second) were captured using an inverted widefield microscope (Zeiss Axio Observer Z1), equipped with a Zeiss Axiocam 506 monochrome camera. Using Zeiss Zen v3.9 software, the diameter of the BA was measured at its widest region, perpendicular to the direction of blood flow, and captured at peak ventricular systolic (maximal BA distension) and diastolic (minimal BA distension) volume (Supplementary Figure 1). BA diameters of compound-treated zebrafish were compared to those of the solvent-treated mutants.

### Bioluminescence assay

*Tg(nppb:Fluc)* or *Tg(nppb:secNluc)* embryos were distributed into white 96-well luminescence plates (Nunc F96 MicroWell, Thermo Scientific) in a total volume of 60 µl E3 medium per well. For preliminary and individual drug assays, one embryo was used per well, whereas six embryos were used for HTDS. 40 µl of luciferase substrate (Steady-Glo or Nano-Glo Luciferase Assay System, Promega, respectively for Fluc and secNluc) was added to each well for an E3:substrate ratio of 3:2. After a 10-minute incubation at room temperature, luminescence was measured using a GloMax Explorer luminometer (Promega).

HTDS was conducted in duplicate by exposure to each compound during two treatment windows, 2-4 dpf and 3-4 dpf, followed by luminescence measurement at 4 dpf. Prior to luminescence readout, embryo morphology was assessed to identify wells showing a beneficial effect (all embryos with a mild phenotype) or evidence of toxicity (all embryos with a severe phenotype or all embryos dead). Wells with toxic effects were excluded from further analyses.

Relative luminescence units (RLU) of each well were log-transformed prior to statistical analysis. Z-scores for compounds were calculated using the formula Z=(x-µ)/σ, where x represents the luminescence value of the compound-treated *fbn3*^-/-^ mutants, and μ represents the mean and σ the standard deviation of the luminescence values of the untreated mutants (negative controls) of the corresponding plate. Compounds were defined as primary hits when both replicates had a Z-score below -1, and as secondary hits when the Z-score of both replicates was below 0.

### Statistical analysis

Continuous data are presented as the mean ± standard error of the mean (SEM) or count (percentage). Group differences of continuous variables were compared using Student’s t-test or Mann-Whitney U test, dependent on normality, while comparisons across multiple groups utilized one-way analysis of variance (ANOVA) with Dunnett’s (comparisons to a control) post-hoc test for multiple comparisons if normally distributed, or Kruskal-Wallis test otherwise. Categorical data (e.g., phenotypic distribution) were analyzed using Fisher’s exact test. Differences between treated and untreated embryos over time were analyzed using Generalized Estimating Equations (GEE) with time as a repeated factor and treatment as the predictor. Dose-dependent trends were assessed using Spearman’s rank correlation. A p-value < 0.05 was considered statistically significant. Statistical analyses were performed using GraphPad Prism version 10.1.2, with the exception of the GEE analysis, which was done in SPSS Statistics version 13.0.0.0.

## RESULTS

### New secreted nanoluciferase reporter line for cardiac stress

To improve the sensitivity of the cardiac stress luciferase reporter system, we generated the novel *Tg(nppb:secNluc)* zebrafish line using a CRISPR/Cas9-based approach. This reporter expresses *nppb* promotor-driven secreted nanoluciferase, which provides a higher dynamic range than Firefly luciferase, and minimizes potential artefacts caused by attenuation of the luminescence signal within cardiac tissue by secretion from cardiomyocytes.

Integration of the reporter construct at the intended *uobl6* safe harbor landing site could, however, not be confirmed by positional PCR using primers spanning the predicted junction between genomic DNA and the transgene. Other PCR-based strategies to map the genomic transgene insertion site were also unsuccessful. Nevertheless, we could amplify the *nppb:secNluc* cassette from genomic DNA of founder zebrafish, showing the expected YFP fluorescence in the lens, indicating that the transgene integrated at a random genomic location. Care was taken to keep experimental zebrafish hemizygous for the transgene allele, and careful analysis did not reveal any adverse effects on transgenic zebrafish viability or development.

To validate the reporter functionally, *Tg(nppb:secNluc)* embryos were treated with compounds known to induce cardiac stress: epinephrine, a β-adrenergic receptor agonist, and 2,3-butanedione monoxime (BDM), a myosin inhibitor that suppresses cardiac contraction. Both epinephrine and BDM treatments increased cardiac stress across most tested concentrations (Supplementary Figure 2), confirming the responsiveness of the *nppb:secNluc* reporter.

### Luciferase-based nppb reporters reveal phenotype-dependent cardiac stress in fbn3 mutants across developmental stages

To assess how *fbn3* loss of function influences activity of the cardiac stress-responsive gene *nppb*, we compared two independent transgenic reporter lines, based on firefly luciferase (*nppb*:*Fluc*) or secreted nanoluciferase (*nppb*:*secNluc*) (Figure 1A and B). *fbn3* mutants expressing each of the luciferases were generated (Figure 1C) and separated into S and M phenotype based on the size of pericardial edema, with severity defined as pericardial edema extending beyond the ventral edge of the yolk sac, as defined in the publication describing the generation and phenotyping of the *fbn3* mutants^15^ (Figure 1D). Luminescence was quantified in heterozygous (*fbn3*^+/-^), mutants with a mild cardiac phenotype (*fbn3*^-/-^ M), and mutants with a severe cardiac phenotype (*fbn3*^-/-^ S) between 4 and 8 dpf (Supplementary Figure 3), and values were expressed as fold change relative to control WT levels (Figure 1F and G).

In both luciferase reporter systems, we found that *fbn3* mutants displayed increased *nppb* promoter activity (Figure 1E), with mutants showing a severe cardiac phenotype consistently showing the highest luminescence, peaking at 7 dpf. *fbn3* mutants with a mild phenotype exhibited elevated signal up to 6 dpf, after which their luminescence declined toward baseline. In contrast, heterozygous larvae remained near wildtype baseline levels throughout the time course.

The two luciferase reporter lines differed in the magnitude of their luminescence responses. While Fluc activity in *fbn3* mutants with a severe phenotype reached just over a 10-fold increase compared with WT levels, the difference in the secNluc line approached a 1000-fold change. This substantially greater dynamic range highlights the enhanced sensitivity and higher enzymatic efficiency of the secreted nanoluciferase reporter in detecting *nppb* regulatory activity.

Together, these results demonstrate that *nppb* expression is robustly induced in *fbn3* mutants, scales with phenotypic severity, and can be sensitively quantified across developmental stages using luciferase-based reporters.

### Testing the effects of TGF-β signaling inhibition on the fbn3^-/-^ cardiovascular phenotype

To assess whether inhibition of TGFβ signaling could modify the phenotype of our *fbn3^-/-^*zebrafish model, we tested two pharmacological inhibitors, RepSox and refametinib, which respectively inhibit the TGF-β type I receptor and the downstream MEK1/2 pathway. Refametinib showed a transient beneficial effect, reducing cardiac stress in *fbn3^-/-^* embryos at 3 dpf. However, this effect was reversed at 4 dpf, where treated mutants displayed increased stress levels compared with controls. Additionally, refametinib consistently increased cardiac stress in WT embryos across all time points (Figure 2A). For the screen of the pericardial phenotype distribution in *fbn3^-/-^* larvae, an additional higher concentration (0.1 µM) was chosen as well to check its efficacy, but neither concentration caused an improvement in the severity of the morphological defects (Figure 2B).

**Figure 2.**
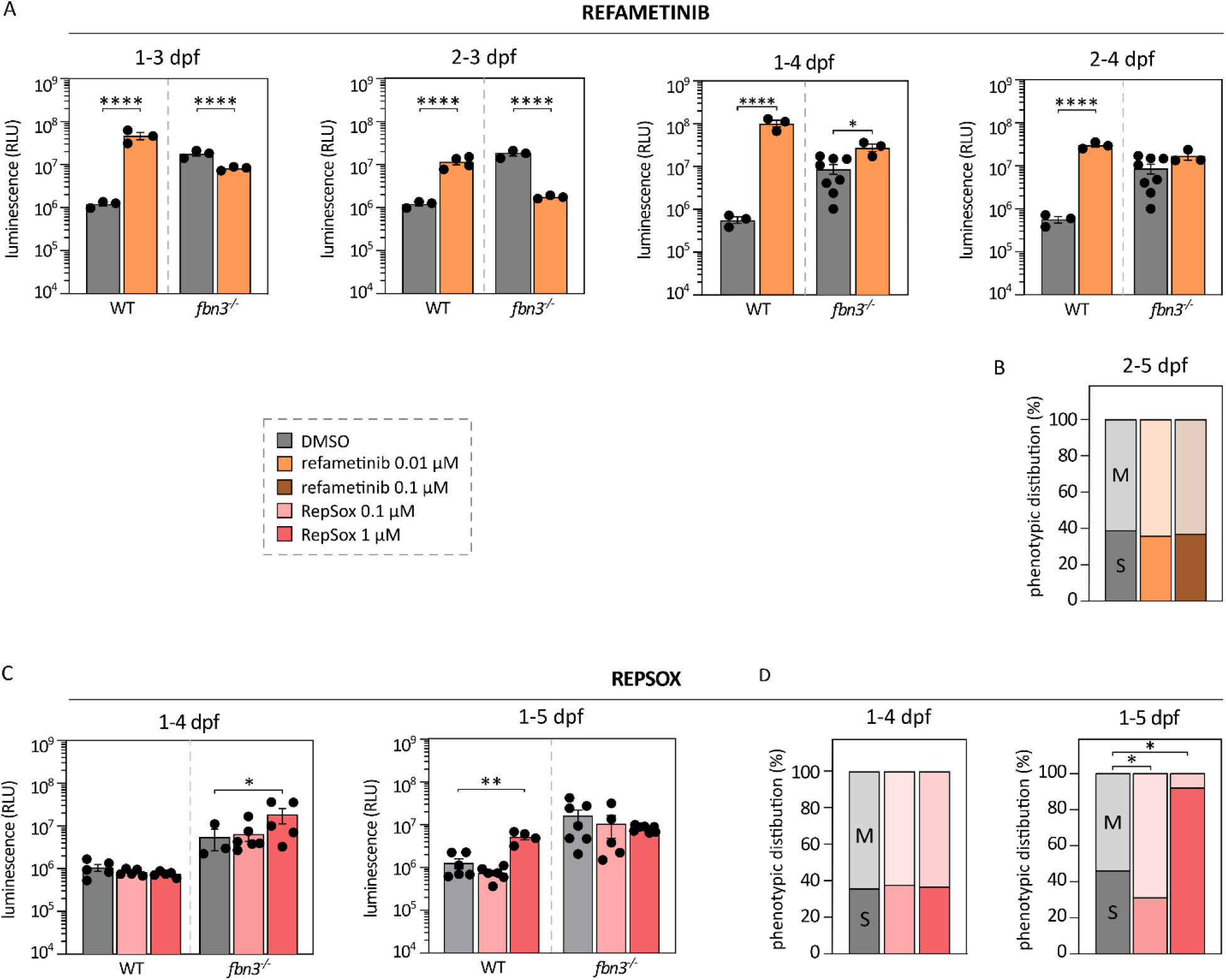
Effects of TGF-β signalling inhibitors on the fbn3^-/-^ zebrafish model of MFS. **(A, B)** *Tg(nppb:secNluc) fbn3^-/-^* zebrafish embryos were treated with the TGF-β signalling inhibitor refametinib (0.01 and 0.1 µM, orange) and **(C, D)** RepSox (0.1 and 1 µM, red), with solvent as a control (DMSO - grey). No beneficial effect on *fbn3^-/-^* phenotype, reflected as lowered luminescence or the shift towards a milder phenotype, was consistently observed for any drug treatment across all incubation times. Displayed time intervals above each of the graphs indicate the start of incubation (first day) and readout (final day). Black dots represent individual larvae. Data are expressed as mean ± SEM. Statistical test analysis: unpaired t-test on log-transformed data (A), Fisher’s exact test (B, D), Kruskal-Wallis test followed by Dunn’s multiple comparisons test on log-transformed data (C). All statistical significance is indicated with asterisks. ****p<0.0001, **p<0.01, *p<0.05. M = mild *fbn3^-/-^* pericardial phenotype, S = severe *fbn3^-/-^* pericardial phenotype.

Based on preliminary screenings, longer incubation periods were applied for RepSox treatment, which was not feasible with refametinib due to toxicity. When treatment was initiated at 1 dpf and assessed at 4 dpf, RepSox had a detrimental effect on cardiac stress in *fbn3^-/-^* larvae as well as in WT larvae when assessed at 5 dpf (Figure 2C). RepSox also caused a pronounced shift toward a more severe pericardial phenotype at 5 dpf (Figure 2D). This indicates that RepSox exacerbates the cardiac phenotype in *fbn3^-/-^* larvae.

### Limited impact of β-adrenergic blockade on fbn3^-/-^ cardiac stress and morphology

To evaluate whether inhibition of the β-adrenergic system, which is the standard treatment strategy in MFS, could modulate the *fbn3^-/-^* phenotype, we initially screened several β-blockers (atenolol, metoprolol, and nebivolol) at their highest tolerable dose without long-term toxicity, using the luminescence-based cardiac stress assay. Among the tested conditions, nebivolol reached the highest tolerated dose and showed the strongest potential, displaying reduced cardiac stress after three days of treatment, although no effect was detected during shorter exposure periods (Figure 3A). When nebivolol was subsequently assessed for longer incubation times and later readout time, this initial trend was not sustained, and no clear reduction in cardiac stress was observed (Figure 3B). Examination of the pericardial phenotype revealed that one specific treatment window produced a transient shift of *fbn3^-/-^* larvae toward a milder phenotype (Figure 3C). However, this apparent improvement was not maintained in a longitudinal assessment with a larger sample size (n = 96). Although treated larvae exhibited a modestly improved phenotype at 3 dpf compared to untreated controls, the phenotype progressively deteriorated over the subsequent two more days of treatment, such that by 5 dpf it was indistinguishable from untreated controls (Figure 3D).

**Figure 3.**
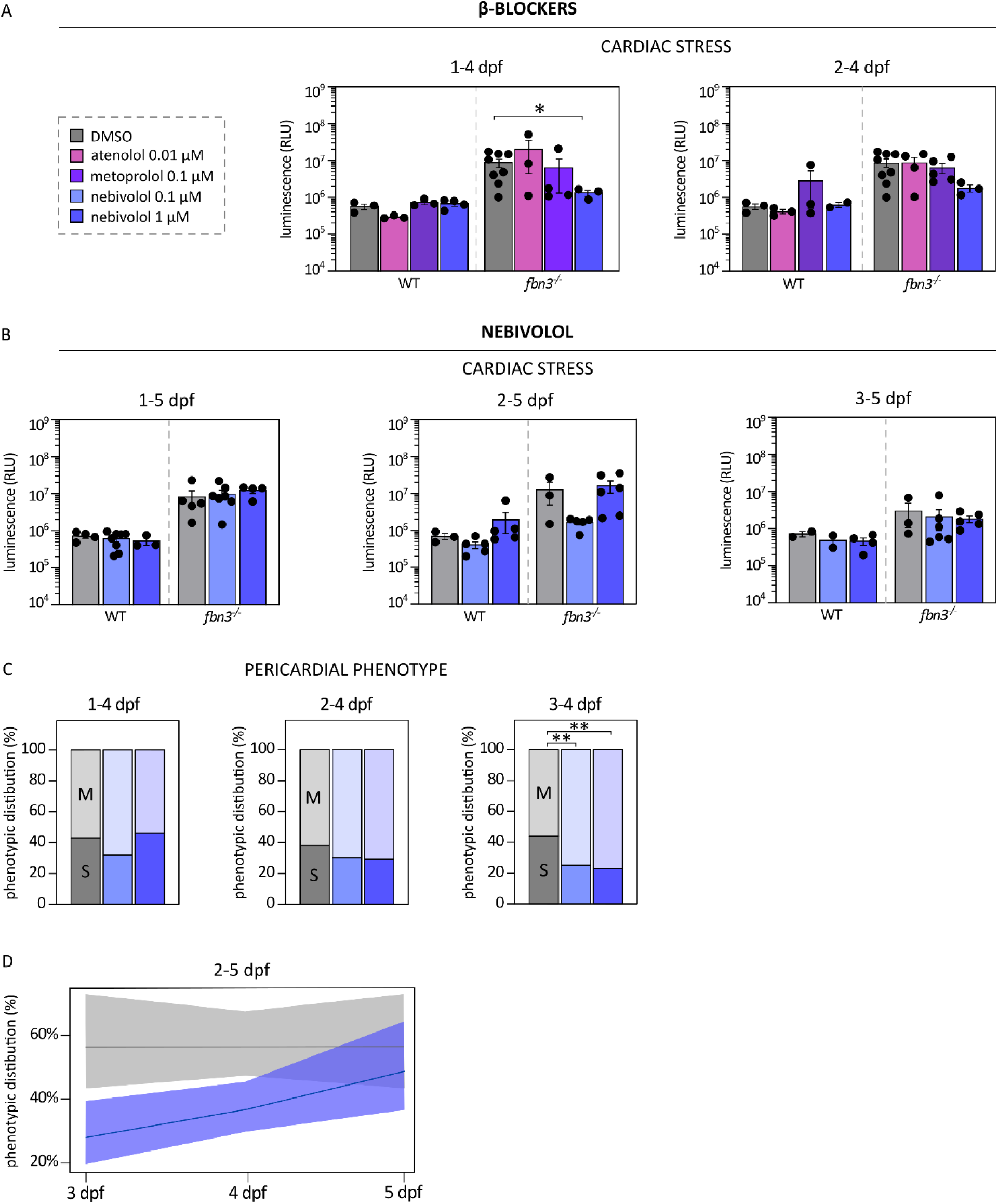
Effects of β-adrenergic receptor blockers on the fbn3^-/-^ zebrafish model of MFS. **(A)** *Tg(nppb:secNluc) fbn3^-/-^* zebrafish embryos were treated with the β-adrenergic receptor blockers atenolol, metoprolol, and nebivolol, and their cardiac stress levels were measured. Based on an observed trend toward reduced cardiac stress with nebivolol, extended treatment experiments were conducted **(B)**; however, this effect was not reproducible. **(C)** Phenotypic distribution data for nebivolol (0.1 and 1 µM), where 1 µM produced a shift toward a milder phenotype. **(D)** The same treatment was therefore repeated longitudinally (scoring individual larvae from 3 dpf until 5 dpf) and at a larger scale (n = 96), but the beneficial effect was not reproduced. Displayed time intervals above each of the graphs indicate the start of incubation (first day) and readout (final day). Black dots represent individual larvae. Data are expressed as mean ± SEM. Statistical test analysis: Kruskal-Wallis test with Dunn’s multiple comparisons test on log-transformed data (A, B), Fisher’s exact test (C), Generalized Estimating Equations test (D). All statistical significance is indicated with asterisks. **p<0.01, *p<0.05. M = mild *fbn3^-/-^*pericardial phenotype, S = severe *fbn3^-/-^* pericardial phenotype.

### Losartan fails to improve the fbn3^-/-^ cardiovascular phenotype

We also evaluated the ARB losartan, a standard therapy prescribed to MFS patients alongside β-blockers. Besides modulation of the renin-angiotensin system, its effects are thought to be mediated in part through attenuation of TGF-β signaling.

Across the five tested concentrations, losartan did not produce beneficial effects on the *fbn3^-/-^* phenotype. No reduction in cardiac stress was observed in *fbn3^-/-^* mutants (Figure 4A), nor was there any improvement in the distribution of pericardial phenotypes at 3, 4, or 5 dpf (Figure 4B). Likewise, losartan treatment failed to alleviate BA dilation, although a Spearman’s rank correlation analysis did reveal a significant trend of smaller BA diameters in maximal distension across increasing concentrations (Figure 4C).

**Figure 4.**
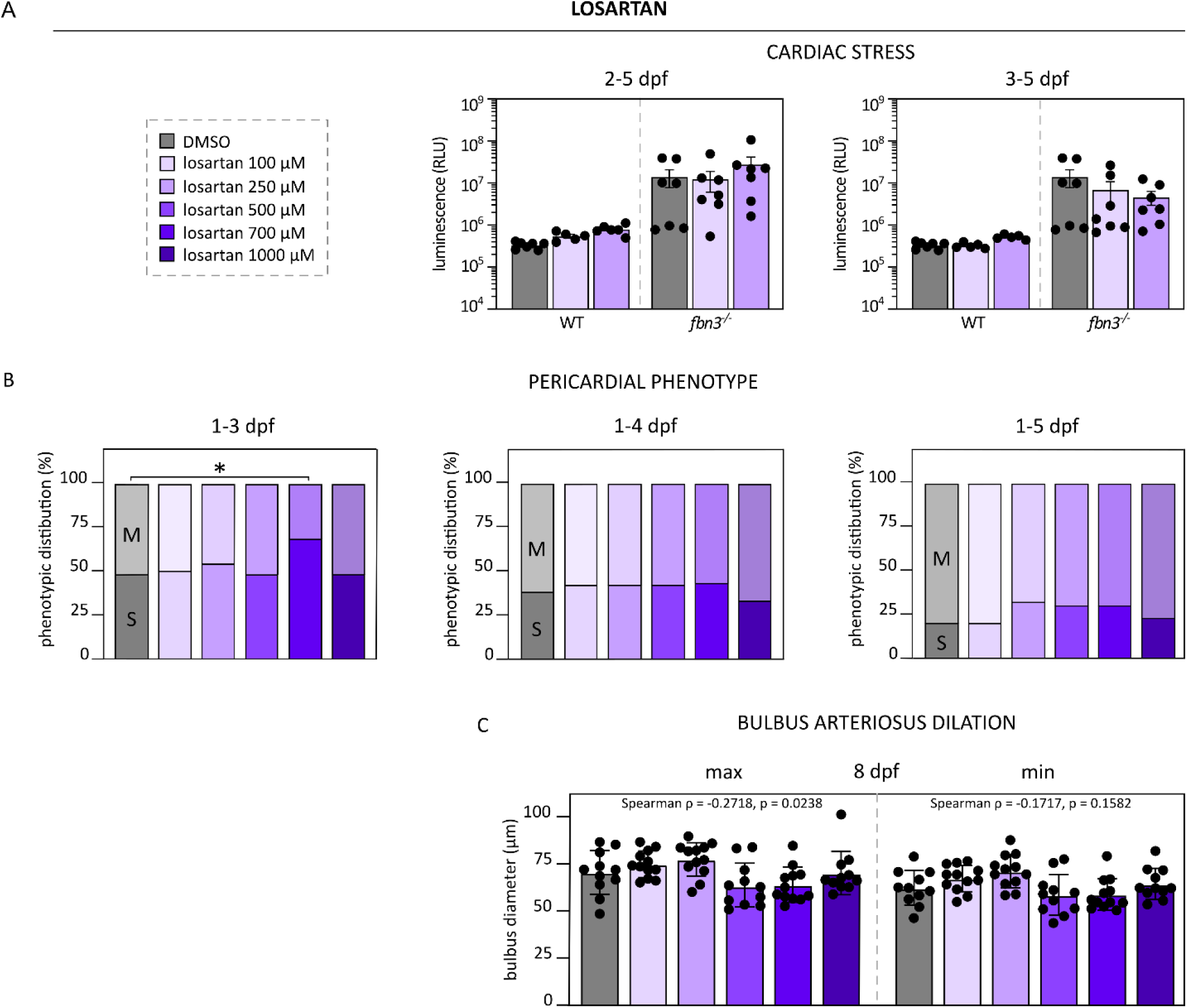
Effects of angiotensin receptor blocker losartan on the fbn3^-/-^ zebrafish model of MFS. *Tg(kdrl:GFP) fbn3^-/-^* zebrafish embryos were treated with the ARB losartan (100, 250, 500, 700, 1000 µM; purple), with solvent as a control (DMSO - grey). There was no beneficial effect on the cardiac stress **(A)**, pericardial phenotype **(B)**, or BA dilation **(C)** of *fbn3^-/-^* phenotype. Displayed time intervals above each of the graphs indicate the start of incubation (first day) and readout (final day). Black dots represent individual larvae. Data are expressed as mean ± SEM. Statistical test analysis: Kruskal-Wallis test with Dunn’s multiple comparisons test (C) on log-transformed data (A), Fisher’s exact test (B), Spearman’s rank correlation test (C). All statistical significance is indicated with asterisks. *p<0.05. M = mild *fbn3^-/-^*pericardial phenotype, S = severe *fbn3^-/-^*pericardial phenotype.

### No improvement of the fbn3^-/-^ phenotype following allopurinol treatment

To evaluate the effect of allopurinol on the *fbn3^-/-^* phenotype, several concentrations were tested across different treatment durations. Allopurinol consistently produced a shift toward a more severe pericardial phenotype across the tested intervals, except the highest concentration, which was comparable to untreated mutants (Figure 5B). Measurements of cardiac stress and BA dilation were comparable across all treatment groups, with no dose-dependent trend, and were indistinguishable from untreated mutants (Figure 5A and C).

**Figure 5.**
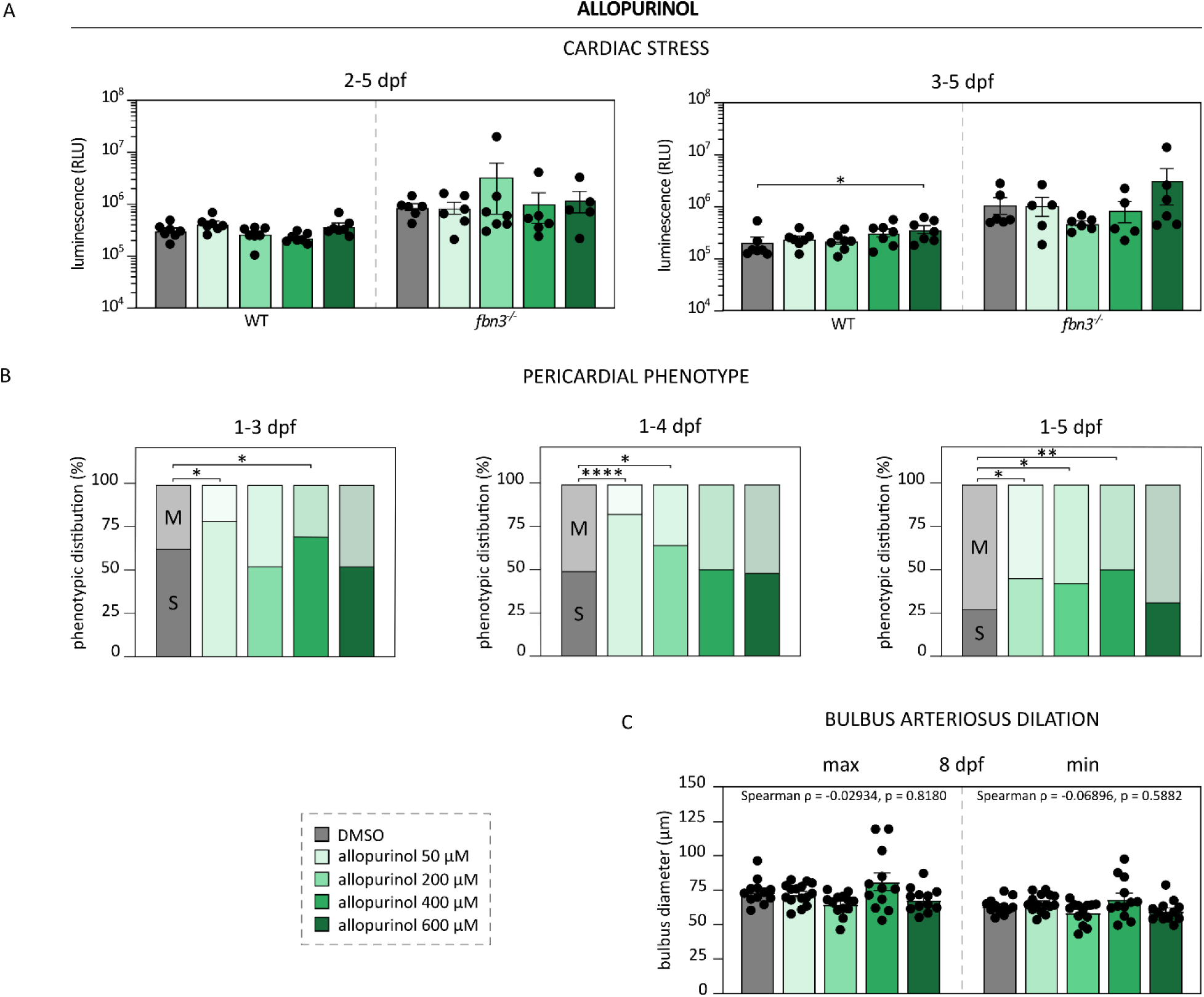
Effects of xanthine oxidoreductase inhibitor allopurinol on the fbn3^-/-^ zebrafish model of MFS. *Tg(kdrl:GFP) fbn3^-/-^* zebrafish embryos were treated with the xanthine oxidoreductase inhibitor allopurinol (50, 200, 400, 600 µM, green), with solvent as a control (DMSO - grey). There was no beneficial effect on the cardiac stress **(A)**, pericardial phenotype **(B)**, or BA dilation **(C)** of *fbn3^-/-^* phenotype. Displayed time intervals above each of the graphs indicate the start of incubation (first day) and readout (final day). Black dots represent individual larvae. Data are expressed as mean ± SEM. Statistical test analysis: Kruskal-Wallis test with Dunn’s multiple comparisons test (C) on log-transformed data (A), Fisher’s exact test (B), Spearman’s rank correlation test (C). All statistical significance is indicated with asterisks. ****p<0.0001, **p<0.01, *p<0.05. M = mild *fbn3^-/-^*pericardial phenotype, S = severe *fbn3^-/-^* pericardial phenotype.

### Optimization of the unbiased high-throughput drug screen in a zebrafish model of MFS

To identify previously unexplored therapeutic candidates capable of modifying the *fbn3^-/-^*phenotype, we performed an unbiased HTDS using a diverse library of compounds with established clinical use. This library was specifically selected to accelerate potential clinical translation, as these compounds have already undergone extensive safety and pharmacokinetic evaluation.

To capture the full phenotypic range of the *fbn3^-/-^*phenotype in one well, we used six embryos per well to ensure a very high likelihood (98.44%) that each well contained at least one severely affected embryo. Assuming an average 50% prevalence of severe phenotypes at 4 dpf^15^, the probability was calculated as 1 − 0.5⁶. This approach accounted for the absence of presorting by phenotypic severity, which was necessary to maintain the high-throughput nature of the screen.

The HTDS was performed in duplicate using two different treatment initiation times. In the first run, compounds were added at 2 dpf, a stage when the earliest signs of endocardial detachment begin to emerge. Treating embryos at this time allowed us to test whether pharmacological intervention could prevent it. In the second run, compounds were added at 3 dpf, when aspects of the phenotype are already more established. This later treatment window enabled us to assess whether the phenotype could still be ameliorated once early pathological changes had already begun.

We selected 4 dpf as the primary luminescence readout time for our HTDS, to be able to capture the most likely time window for treatment. In our preliminary experiments, we found that incubating zebrafish embryos from 2-5 dpf in a 96-well plate severely affected survival even without drug treatment. In addition, by 5 dpf, the phenotypic distribution becomes more constrained: fewer embryos remain in the severe category, and those that are severe typically show advanced pathology, including pronounced pericardial and yolk sac edema and increasing mortality^15^, which could negatively skew the readouts. We therefore chose to perform the HTDS with drug incubation times from 2-4 dpf and 3-4 dpf. Using these two exposure regimens allowed us to evaluate both preventive and early rescue effects, and to measure effects over different developmental time windows, ensuring that the screen captured the full therapeutic potential of each compound across critical stages of disease development.

### Unbiased high-throughput drug screen identifies no reproducible beneficial candidates

After exposure of *fbn3^-/-^* embryos on a *Tg(nppb:secNluc)* background to the clinically approved drug library either from 2-4 or 3-4 dpf, survival and zebrafish morphology were checked using brightfield microscopy observation. Afterwards, a luciferase assay was performed, and luminescence was quantified relative to untreated mutants, which were included on each plate as a negative control. Z-scores were calculated for both screening rounds, and selected hits were further validated (Figure 6).

**Figure 6.**
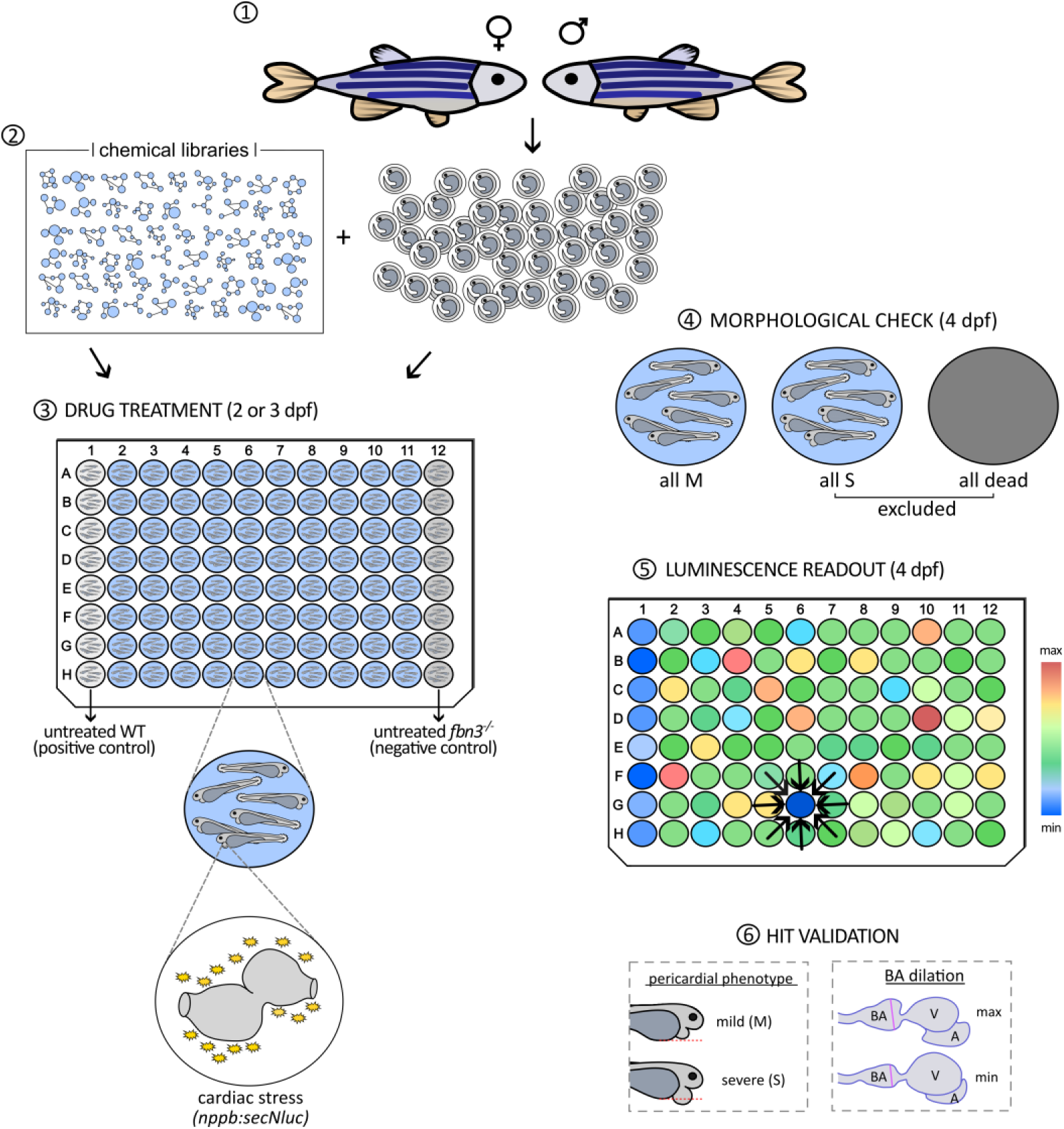
Workflow of a high-throughput drug screen on fbn3^-/-^ zebrafish model of MFS. **(1)** Adult *Tg(nppb:secNluc) fbn3^-/-^* zebrafish are crossed to generate a high number of embryos. **(2)** A diverse chemical library of over 1500 compounds, previously approved or developed for other applications, was screened. **(3)** At 2 dpf (3 dpf for the duplicate screen), embryos were distributed to 96-well plates containing individual compounds at a final concentration of 10 µM. Positive control (WT) and negative control (*fbn3^-/-^*) were treated with a solvent only (DMSO). **(4)** At 4 dpf, each well was examined for morphology to identify wells showing a beneficial effect (all M - all embryos displaying a mild phenotype) or signs of toxicity (all S - all embryos displaying a severe phenotype, or all embryos dead). Wells with toxic effects were excluded from further analyses. **(5)** The main readout, luminescence from a cardiac stress reporter (*nppb:secNluc*), was quantified to identify compounds that reduce stress-induced signal intensity. **(6)** Candidate hits were further validated through assessment of cardiac phenotypes, including pericardial phenotype (expressed as the percentage embryos with severe phenotype) and BA dilation, across a range of drug concentrations and incubation timings. M = mild *fbn3^-/-^*pericardial phenotype, S = severe *fbn3^-/-^*pericardial phenotype.

Across both duplicates, 16.5% of the tested compounds produced Z-scores below 0, classified as secondary hits, indicating reduced cardiac stress relative to untreated mutants (Figure 7A). The most represented therapeutic categories within this group included antibacterial, analgesic/anti-inflammatory, antipsychotic, antiepileptic, antiparasitic, antihyperlipidemic, and central nervous system modulators, as listed in Figure 7A. A smaller subset, representing 1.3% of all screened compounds, showed Z-scores below -1 (Figure 7B), classified as primary hits. From both primary and secondary hits, 12 compounds were selected for further validation in more detail. Each compound was tested across six concentrations (1 nM - 100 μM) and three treatment windows (1-6 dpf, 2-6 dpf, and 3-6 dpf). Pericardial phenotype was assessed between 3 and 5 dpf, followed by analysis of BA dilation at 6 dpf in the same group of treated embryos. Phenotypic scoring criteria for pericardial severity and BA diameter were standardized (Figure 7D). For each concentration and treatment window, the resulting pericardial phenotype distribution and BA dilation measurements were recorded and used to generate a cardiotoxicity-cardioprotection heatmap summarizing treatment effects across the entire validation panel (Figure 7C).

**Figure 7.**
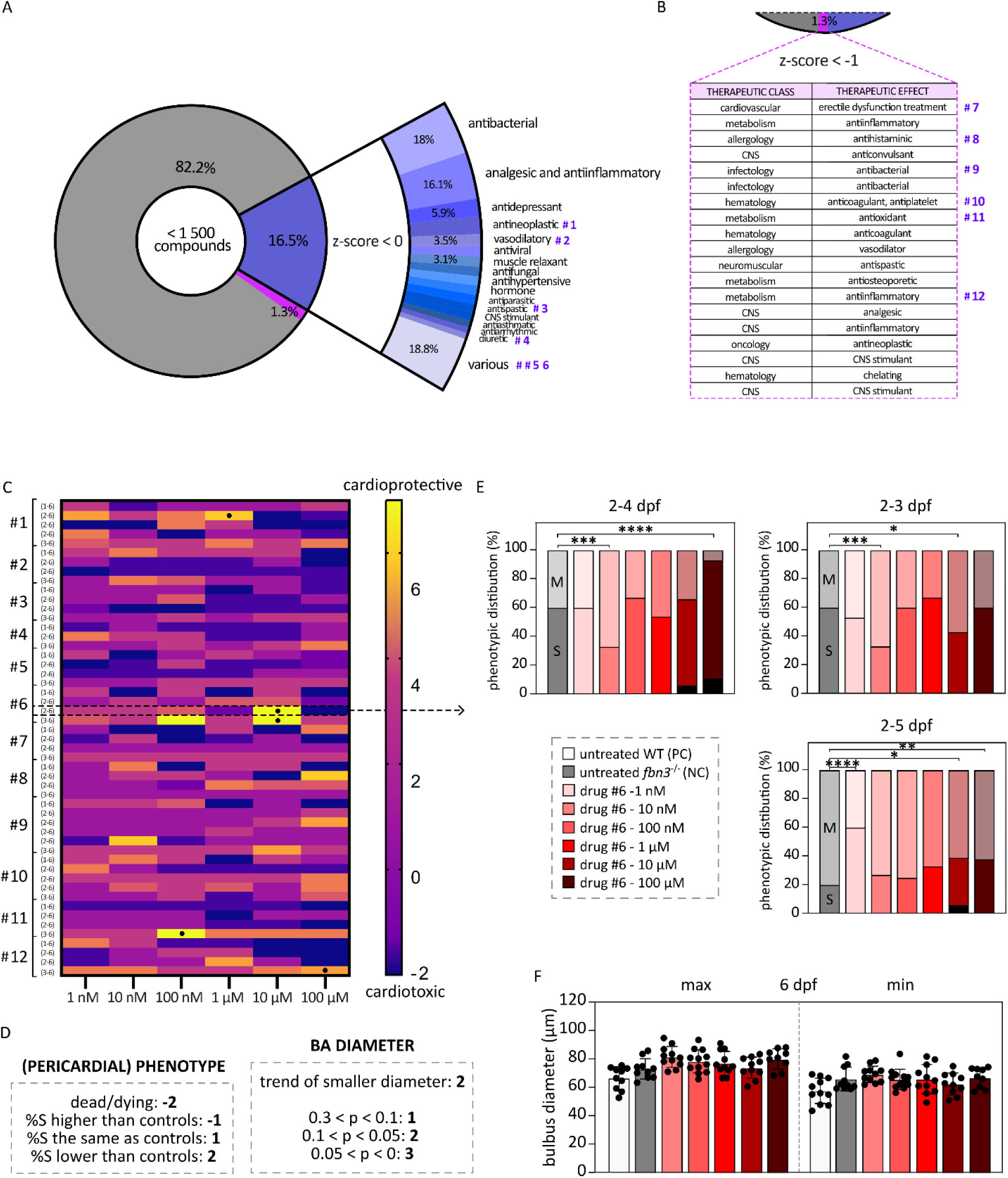
High-throughput drug discovery and follow-up hit validation. Distribution of >1500 screened compounds by z-score, highlighting the percentages classified as secondary hits with a z-score < 0 (**A**, blue) or primary hits with a z-score < –1 (**B**, pink), together with their most common therapeutic categories. The **#** labels denote the hit compounds selected for further validation. (**C**) Heatmap showing validation of 12 hits across six drug concentrations (1 nM - 100 μM), ranging from the lowest cardiotoxic scores (blue) to the highest cardioprotective scores (yellow). Each row represents a replicate; numbers in brackets indicate the treatment time windows. Conditions that were further validated in larger sample sizes are indicated with a dot (•). Dashed lines highlight the representative compound analyzed in panels E and F. (**D**) Scoring system used to generate the heatmap, including pericardial phenotypes (percentage of severe embryos per group, %S, relative to untreated mutants) and BA diameter. (**E**) Phenotypic distribution of *fbn3^-/-^* embryos treated with compound #6 from 2 dpf to 6 dpf (treatment window marked by the dashed line in panel C), compared with untreated mutants (grey). The percentage of dead embryos is shown in black. (**F**) BA diameter measurements at 6 dpf for embryos treated with compound #6 across the concentration series. Controls include untreated WT (positive control, PC) and untreated mutants (negative control, NC). Displayed time intervals above each of the graphs indicate the start of incubation (first day) and readout (final day). Black dots in panel F represent individual larvae. Data are expressed as mean ± SEM. Statistical test analysis: Fisher’s exact test (E), Kruskal-Wallis test with Dunn’s multiple comparisons test (F). All statistical significance is indicated with asterisks. ****p<0.0001, ***p<0.001, **p<0.01, *p<0.05. M = mild pericardial phenotype, S = severe pericardial phenotype.

Representative examples of phenotype distribution shifts for one candidate compound are shown in Figure 7E, and corresponding BA diameter measurements at maximal and minimal distension at 6 dpf are shown in Figure 7F. Compounds that appeared most promising in the initial validation were subsequently retested at larger sample sizes for both phenotype distribution and BA morphology. However, none of the treatments demonstrated reproducible effects in the expanded validation assays.

## DISCUSSION

Pharmacological management of the cardiovascular manifestations of MFS has remained relatively unchanged over the years. β-blockers have long been the standard of care, primarily aimed at reducing hemodynamic stress on the weakened aortic wall. The introduction of ARBs, particularly losartan, generated considerable enthusiasm following striking preclinical findings in which treated MFS mice exhibited near-normalization of the aortic phenotype, becoming virtually indistinguishable from WT controls^26^. However, the benefits observed in murine models were not fully replicated in humans^27–29^. Clinical studies also failed to demonstrate a clear superiority of losartan over β-blockers^30–32^, with subsequent analyses suggesting that early combination therapy may provide greater attenuation of aortic enlargement than either treatment alone, particularly with long-term use^13,33^. Nevertheless, these pharmacological strategies primarily slow disease progression and delay the need for surgical intervention; they do not eliminate the risk of aortic dissection and rupture, which still occurs even under close medical management^13,29^.

It is concerning that, despite more than a century of research, no medical treatments are available to halt MFS progression or fully prevent its most severe complications. Therefore, there is a strong need for novel animal models that can facilitate the discovery and evaluation of new therapeutic strategies. In the current study, we describe a new zebrafish model that provides a platform for both targeted and unbiased screening of potential treatments for the cardiovascular manifestations of MFS.

### Elevated cardiac stress in a fbn3^-/-^ zebrafish MFS model

We generated a novel *Tg(nppb:secNluc)* reporter line, which shows a significantly larger dynamic range than the previously published firefly luciferase-based *Tg(nppb:Fluc)* reporter line. Despite the fact that we could not confirm insertion of the transgene into the intended genomic safe harbor locus, we proceeded working with the new reporter line as functional testing of the luminescence readout was very successful. We assume that the transgenic cassette was inserted randomly into the genome, similar to how the *Tg(nppb:Fluc)* line was originally generated. *fbn3^-/-^*mutants exhibited increased cardiac stress, evidenced by up to a 1000-fold rise in luminescence in the *secNluc* reporter line (compared to a 10-fold increase in the Fluc reporter line), peaking at 7 dpf. This time point coincides with the onset of mortality in severely affected mutants due to the obstruction of blood flow caused by endocardial detachment, as we previously reported^15^. In contrast, mildly affected mutants showed elevated signal until 6 dpf, after which luminescence declined toward baseline, suggesting progressive cardiac recovery. These zebrafish survive normally to adulthood with preserved cardiac function despite structural valve defects, as we previously reported.

Owing to its greater dynamic range, enhanced sensitivity, and higher enzymatic efficiency compared to Fluc, the secNluc transgenic line was selected for subsequent drug screening.

### Targeting TGF-β signaling in a zebrafish model of MFS

Early preclinical studies of MFS have revealed both protective and pathogenic roles of TGF-β signaling, depending on developmental stage and disease context, underscoring its complexity as a therapeutic target^34,35^. In this study, we investigated refametinib, a highly selective ERK1/2 inhibitor that modulates the noncanonical TGF-β pathway, and RepSox, an inhibitor of the TGF-β type I receptor. Refametinib has previously been shown to significantly reduce aortic root growth in the *Fbn1^C1041G/+^* mouse model^36,37^, whereas RepSox induced severe dilation of the outflow tract in a zebrafish knockout model^38^.

In our experiments, both compounds exhibited mixed effects on the outcome of *fbn3^-/-^*larvae. Refametinib showed a beneficial effect on cardiac stress in 3 dpf *fbn3^-/-^* larvae after 1 or 2 days of drug exposure, but this effect was not seen at later stages. This may indicate a transient developmental window in which the mutant phenotype is sensitive to MEK/ERK pathway inhibition, while later stages may involve disease progression or compensatory mechanisms that reduce treatment responsiveness. However, refametinib consistently increased cardiac stress at all time points in WT larvae, indicating that its effects are highly stage- and genotype-dependent and that MEK/ERK inhibition might interfere with normal cardiac development and/or homeostasis at these stages. In contrast, treatment with 1 µM RepSox increased cardiac stress in 4 dpf *fbn3^-/-^* larvae and increased the proportion of severely affected *fbn3^-/-^* embryos at 5 dpf. These data are in line with previous results obtained in mouse models, highlighting the context-dependence of modulation of TGF-β signaling in the setting of MFS.

### Standard MFS therapies have no strong effect on the fbn3^/-^ phenotype

To evaluate the effects of commonly prescribed medication for MFS in our zebrafish model, we tested three β-blockers: the second-generation β₁-selective blockers atenolol and metoprolol, and the third-generation agent nebivolol, which combines high β₁ receptor specificity with endothelium-dependent vasodilatory properties. Among these, only nebivolol showed a modest beneficial effect on the phenotype of *fbn3^-/-^* larvae. However, this effect was not reproducible in larger cohorts or across different treatment windows.

The ARB losartan, another standard therapy in MFS, only led to a mild dose-dependent improvement in the BA dilation of *fbn3^-/-^* larvae, but no significant improvements were seen in the cardiac stress readout or in the pericardial edema severity. These findings are consistent with clinical observations indicating that current mediation can slow the progression of aortic dilation and delay the need for prophylactic surgery, yet fails to eliminate the risk of aortic dissection and rupture, which persists even under careful medical management^13^.

Recent preclinical data showed that allopurinol could prevent or halt the progression of aortic aneurysm in a mouse model of MFS^39^, suggesting that this antioxidant compound might represent a new therapeutic strategy for the treatment of MFS. Nevertheless, administration of allopurinol did not improve pericardial phenotype severity, cardiac stress, or BA dilation in *fbn3^-/-^*larvae.

The absence of a strong effect of losartan and allopurinol in our zebrafish model may reflect limited conversion of these compounds into their active metabolites, EXP3174 and oxypurinol, respectively. This possibility would need to be addressed in future validation experiments, for example, by assessing compound metabolism or directly testing the active metabolites. Alternatively, the lack of a clear beneficial effect may indicate that, under the conditions tested, these compounds do not sufficiently modulate the key pathophysiological mechanisms driving the cardiovascular manifestations of MFS in this model. Together, these findings highlight the need for further pharmacological validation and support the exploration of novel therapeutic strategies.

### Unbiased high-throughput drug screen in a zebrafish model of MFS

The proportion of primary hits (1.3%) is expected for an unbiased screen of this scale and diversity, and in fact supports the robustness of the assay, as higher hit rates in such contexts frequently reflect nonspecific activity or assay interference rather than true biological effects. In this context, we also included secondary hits, compounds that produced reproducible, though more modest, phenotypic rescue, as even partial effects can provide meaningful biological insight and may reflect dose-dependent, context-specific, or early-stage therapeutic potential. Additionally, weaker hits may reflect suboptimal pharmacokinetics and/or pharmacodynamics, limited bioavailability, or context-dependent activity, and could show enhanced efficacy upon further lead optimization or in combination therapies.

The largest category of hits consisted of antibacterial compounds. While it may seem counterintuitive that agents targeting bacterial replication or morphology could ameliorate aortic pathology, it is conceivable that these compounds have additional biological activity. Interestingly, similar unexpected findings have previously been reported in the MFS field; for example, doxycycline, a typical tetracycline antibiotic, has been shown to inhibit MMP activity and has been used in preclinical mouse studies^40,41^. Anti-inflammatory compounds were also common among the secondary hits, which is interesting, as growing evidence implicates chronic inflammation in the initiation and progression of aortic disease^42,43^. Components of both the innate and adaptive immune systems are commonly present within the aneurysmal wall, and in the *fbn3^-/-^* model, we observed upregulation of the complement system alongside increased neutrophil recruitment^15^.

While many of the identified hits exhibited these therapeutic effects already associated with MFS pathology, the strength of an unbiased HTDS lies in its ability to uncover previously unrecognized disease mechanisms. We therefore avoided restricting our selection to compounds aligned with existing, and still incomplete, knowledge. Instead, we selected 12 hits associated with different signaling mechanisms in an unbiased manner for further validation across multiple time windows and drug concentrations. However, systematic scoring and integration of these results revealed notable variability in treatment responses for most compounds, even under comparable experimental conditions, as reflected in the combined heatmap.

This variability likely reflects underlying biological complexity rather than merely experimental inconsistency. Differences in developmental stage at the time of treatment may significantly influence drug responsiveness, as the cardiac phenotype and its associated pathways evolve over time. In addition, the effects of certain compounds may depend on precise dosing, with narrow or non-linear response windows leading to variable outcomes across concentrations. Biological heterogeneity between embryos likely contributes to this variability. This is particularly evident in the *fbn3^-/-^* line, where we have broadly classified mutants as having mild or severe phenotypes for analysis purposes, yet a subset of mutants (approximately 30 %) seems to recover between 3-5 dpf, reverting from severe to mild phenotype^15^. These differences in baseline phenotype severity or physiological state may further contribute to differences in treatment efficacy. Moreover, some compounds may act on interconnected or compensatory pathways, resulting in context-dependent or partial phenotypic rescue.

So far, no definitive hit has yet been shown to consistently ameliorate the *fbn3^-/-^*phenotype, underscoring the complexity of the disease processes. In ongoing studies, we are extending the chemical space of our HTDS by screening larger, more diverse compound libraries to enhance the chances of identifying novel therapeutic candidates.

### Study limitations

This study has several limitations. First, although the *fbn3^-/-^* zebrafish model captures key features of Marfan-associated cardiovascular pathology, it may not fully recapitulate the complexity of human disease, particularly aspects that emerge later in life. These early-developmental assays, therefore, might miss therapeutic benefits that manifest only at later stages or require long-term remodeling of cardiovascular tissues. Second, phenotypic variability within the mutant line, ranging from mild to severe presentations, introduces biological heterogeneity that can obscure drug effects and contribute to variable treatment responses. We have also observed some limited batch effects, with different clutches (embryos collected from different mating pairs) showing different proportions of the *fbn3^-/-^* pericardial phenotype, ranging from approximately 20 to 60% of offspring at 4 dpf showing the more severe, lethal phenotype (Supplementary Figure 4). By mixing embryos from different clutches before starting experiments, particularly in the HTDS setup, we are able to mitigate this confounding factor. Nevertheless, for smaller validation experiments, this is an inherent limitation of the *fbn3* mutant model.

Additionally, differences in drug absorption, metabolism, and bioavailability between zebrafish and mammals may limit the translational relevance of certain findings. Finally, while the HTDS provides a broad chemical coverage, the reliance on single-agent assays and limited dosing windows may overlook compounds that act synergistically or require precise temporal targeting. Nevertheless, this study lays important groundwork for identifying new therapeutic strategies beyond standard MFS care, highlighting the potential of zebrafish-based screening to guide future investigations in complementary models.

## CONCLUSION

This study highlights the urgent need for more effective therapeutic strategies for MFS, as current treatments slow but do not stop disease progression or fully prevent life-threatening complications. Our *fbn3^-/-^* zebrafish model provides a promising platform for early, quantifiable assessment of cardiac and vascular phenotypes, which enables systematic evaluation of potential therapies. Neither established MFS medications nor targeted modulation of TGF-β signaling produced consistent or meaningful rescue in this model, underscoring the limitations of existing approaches. Although our initial unbiased chemical screen did not yet yield a definitive therapeutic candidate, it revealed potential biologically relevant pathways and emphasized the complexity of disease mechanisms. Additionally, combination therapies could be a relevant option as well, as the complex nature of MFS may require more than one treatment target. Finally, expanding this screening effort with larger and more diverse compound libraries holds promise for identifying novel interventions capable of altering the course of MFS and improving patient outcomes.

## LIST OF ABBREVIATIONS

ARB: angiotensin II receptor blockers
BA: bulbus arteriosus
BDM: 2,3-butanedione monoxime
BMP: bone morphogenetic proteins
DMSO: dimethyl sulfoxide
dpf: days post-fertilization
ECM: extracellular matrix
FBN1: fibrillin-1
fbn3^-/-^: fibrillin-3
Fluc: Firefly luciferase
GEE: Generalized Estimating Equations
HTDS: high-throughput drug screen
M: mild (*fbn3^-/-^*phenotype)
MFS: Marfan syndrome
ORI: origin of replication
RLU: relative luminescence units
S: severe (*fbn3^-/-^*phenotype)
secNluc: secreted nanoluciferase
SEM: standard error of the mean
TGF-β: transforming growth factor-β
VSMC: vascular smooth muscle cell
WT: wild-type
YFP: yellow fluorescent protein

## DECLARATIONS

### ETHICS APPROVAL AND CONSENT TO PARTICIPATE

All experiments were approved by the Animal Ethics Committee of the Ghent University Faculty of Medicine and Health Sciences (ECD 19-16K) and conform to the guidelines from Directive 2010/63/EU of the European Parliament on the protection of animals used for scientific purposes.

### CONSENT FOR PUBLICATION

Not applicable.

### AVAILABILITY OF DATA AND MATERIALS

All data generated or analysed during this study are included in this published article and its supplementary information files.

### AUTHORS’ CONTRIBUTIONS

MH and PS conceptualized the study and designed the methodology. MHR, LM, ED, JW, EV, IR collected the data, while MHR led the formal data analysis and wrote the initial draft. LC, KDR, LM, ED, JW, EV, IR, IRR, GE, JDB and PS reviewed the methodology, results of the study, and overall the manuscript. JDB and PS supervised and approved the work. All authors have read and approved the manuscript.

## ACKNOWLEDGEMENTS

We thank the Zebrafish Facility Ghent Core, and in particular Karen Vermeulen, for the excellent care of the zebrafish. We are also grateful to Dr. Calum MacRae for providing the plasmid and transgenic zebrafish line.

