## Supplementary Appendix for "Integrated luminescence and phenotypic profiling for drug discovery in a zebrafish model of Marfan syndrome"

***Supplementary Table 1. Primer pairs used for nppb:secNluc reporter construction.***

| **GENE** | **FORWARD PRIMER** | **REVERSE PRIMER** |
| --- | --- | --- |
| *nppb* promoter | CCTGCATCAGAGTATGGTG | GTCTCCTGATATACTTTTTTTTTTTTAAATC |
| *secNluc* | ATGAACTCCTTCTCCACAAG | GGGTTGAAGGCTCTCAAG |
| *α-crystallin:YFP+ORI+AmpR* | ATAATTCACTGGCCGTCG | CACTTTTCGGGGAAATGTG |

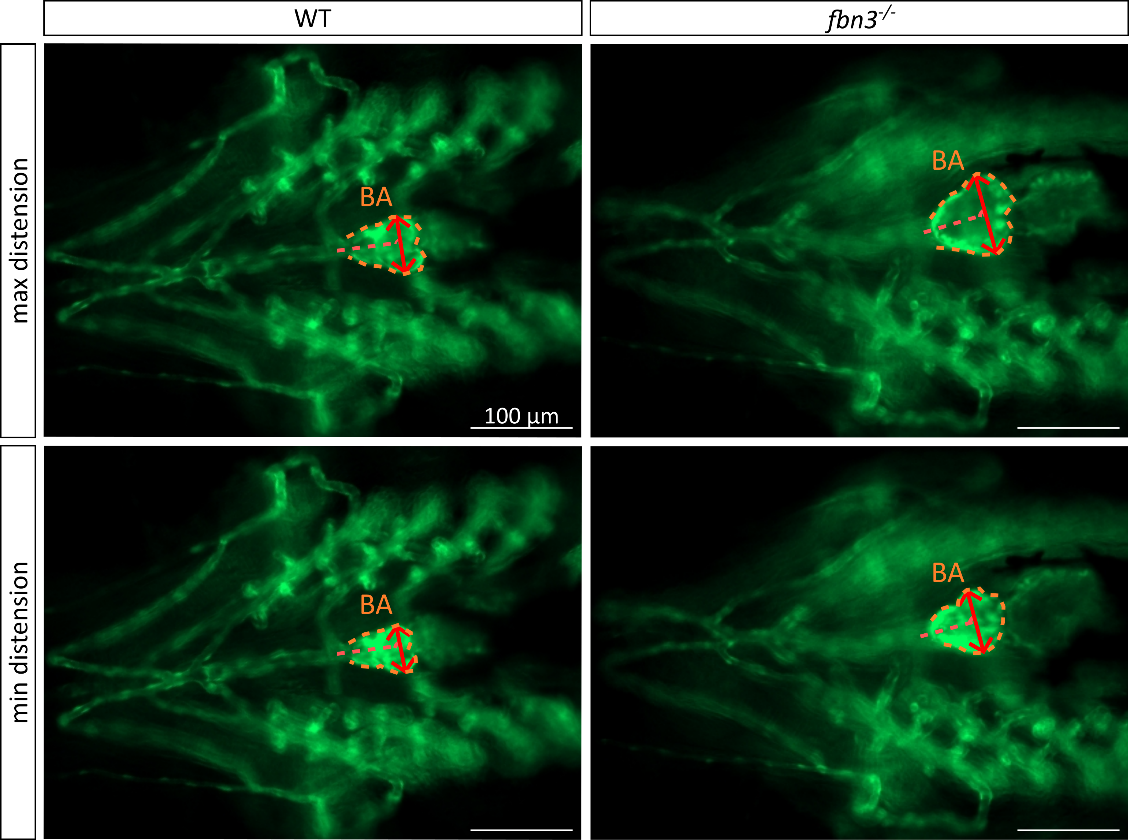

***Supplementary Figure 1 - BA morphology and distension dynamics in WT and fbn3^-/-^ zebrafish larvae.***

Representative live imaging of the BA imaging of *Tg(kdrl:GFP)* 6 dpf larvae, acquired from a ventral perspective over a full cardiac cycle. The BA is outlined (orange dashed line), and the measurement axis (red line) marks the plane along which BA diameter was quantified. Measurements were taken at the widest region of the BA, perpendicular to the blood flow direction, and captured at peak ventricular systolic (max distension) and diastolic (min distension) phases. Scale bar: 100 μm.

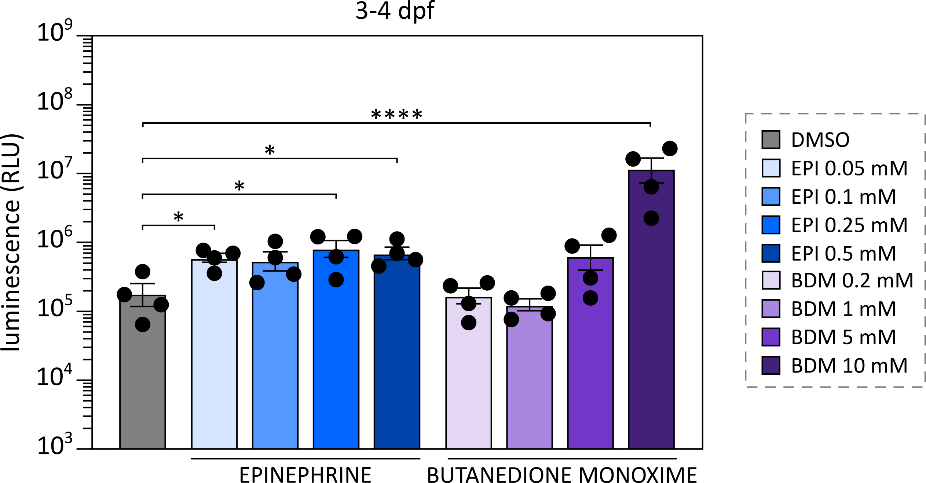

***Supplementary Figure 2 – Validation of the new Tg(nppb:secNluc) reporter line using known cardiac stressors.***

*Tg(nppb:secNluc)* WT zebrafish embryos were treated with epinephrine (EPI) (0.05, 0.1, 0.2, 0.5 mM, blue) and butanedione monoxime (BDM) (0.2, 1, 5, 10 mM), purple), with solvent as a control (DMSO - grey). The graph shows relative luminescence units (RLU) of individual larvae (black dots). There is an increase in cardiac stress in response to most tested epinephrine concentrations, as well as a dose‑dependent increase following BDM treatment. Displayed time interval indicates the start of incubation (first day) and readout (final day). Data are expressed as mean ± SEM. Statistical test analysis: one-way ANOVA followed by Dunnett's multiple comparisons test on log- transformed data. All statistical significance is indicated with asterisks. ****p<0.0001, *p<0.05.

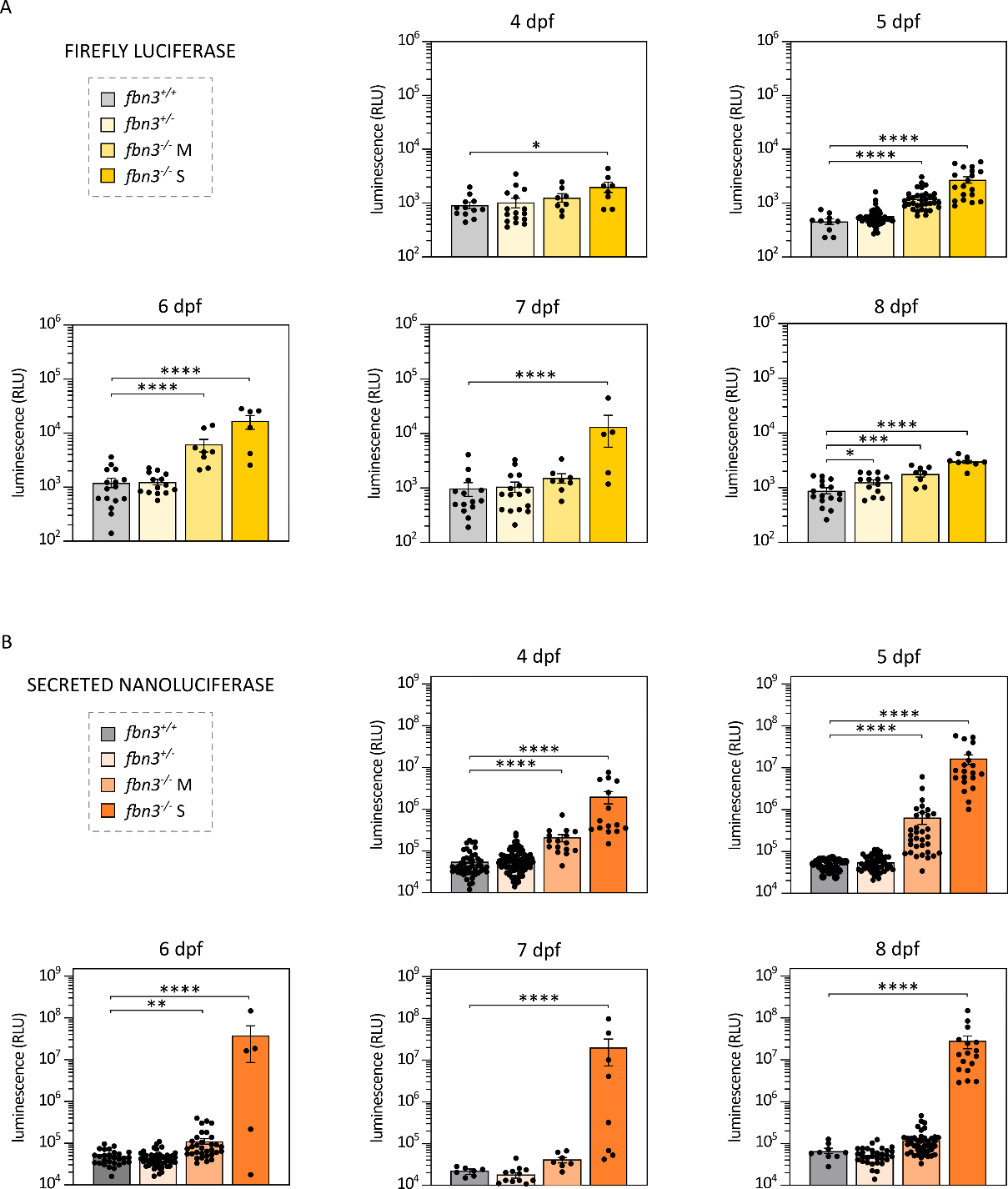

***Supplementary Figure 3 - Luciferase-based‑ quantification of cardiac stress during early development in fbn3^‑/‑^ mutant zebrafish using Fluc and secNluc reporters.***

**(A)** Firefly luciferase-based cardiac stress reporter signal from *Tg(nppb:Fluc)* transgenic zebrafish line measured in four genotypes across 4-8 days post‑fertilization (dpf). Each panel shows relative luminescence units (RLU) of individual larvae (black dots), illustrating developmental changes in reporter activation and genotype‑dependent differences in cardiac stress. The highest luminescence levels are consistently detected in the *fbn3^‑/‑^* zebrafish, particularly within the severe subgroup. **(B)** Secreted nanoluciferase-based cardiac stress reporter signal from *Tg(nppb:secNluc)* transgenic zebrafish line measured in the same genotypes and developmental stages. Compared with the Firefly reporter, the secNluc system displays higher dynamic range and more pronounced genotype‑dependent differences in cardiac stress. Statistical test: one-way ANOVA followed by Dunnett’s multiple comparisons test on log-transformed data. All statistical significance is indicated with asterisks. ****p<0.0001, ***p<0.001, **p<0.01, *p<0.05. M = mild pericardial phenotype, S = severe pericardial phenotype.

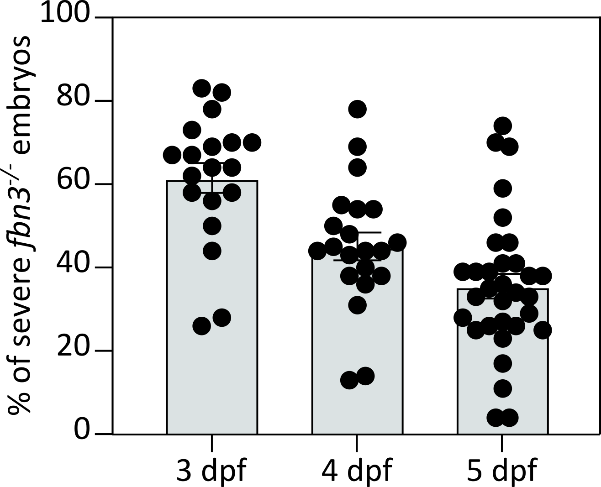

***Supplementary Figure 4 – Clutch-dependent variability in phenotypic severity of fbn3^-/-^ embryos.***
The percentage of severely affected *fbn3^-/-^* embryos was quantified for independent clutches at 3, 4, and 5 dpf, each dot representing a single clutch (all embryos from a single mating pair). The graph presents a new representation of the data originally reported in De Rycke *et al*.^1^ This visualization highlights the separate data points to demonstrate the inter‑clutch variability in phenotypic severity and its progression over time, which was not captured in the previous analysis.

***REFERENCES***

1.	De Rycke K, Horvat M, Caboor L, et al. Systematic Disruption of Zebrafish Fibrillin Genes Identifies a Translational Zebrafish Model for Marfan Syndrome. *JACC Basic Transl Sci*. 2026;11(5):101543. doi:10.1016/j.jacbts.2026.101543
